# Role of pharmacists and community pharmacies in screening, knowledge and awareness investigation about diabetes mellitus type 2 of Jordanian people visiting community pharmacies

**DOI:** 10.1101/2022.11.29.518336

**Authors:** Anas Khaleel, Mona Abu-Asal, Abdullah Bassam Zakariea, Rowan Alejielat, Anas Z. Al-Nweiran

## Abstract

**Background:** The problem is that approximately half of people with diabetes are unaware that they have the disease. Because there are few signs or symptoms in the early stages of diabetes, unnoticed symptoms will persist until diabetic complications appear just before D.M. is diagnosed. Diabetes is increasing exponentially worldwide, particularly in Africa and the Middle East. This study aims to determine Jordanians’ awareness of type 2 diabetes among those who visit community pharmacies in Amman, Jordan, as well as clarify the role of community pharmacies in T2DM screening.

**Methods:** The design was based on participants who visited community pharmacies in Amman, Jordan, in 2021. The personal contact interview questionnaire collected demographic information, geographic location, educational attainment, and insurance status. In addition, we created 15 knowledge questions. The study included 305 participants. Descriptive and regression analyses were deployed by using SPSS,

**Results:** A significant relationship between the type of medical degree and knowledge of risk factors for type 2 diabetes mellitus was confirmed in this study (p <0.012). Some subjects scored slightly higher than others (n = 175; 57.4% of participants scored above 7, compared to n = 130; 42.6% scored below 7). Although 50.5% of the participants (n = 154) held a bachelor’s, master’s, or doctorate, these degrees did not improve the participants’ knowledge levels. The association was tested using chi-square analysis, but no significance was found.

**Conclusions:** Random visitors to Jordanian community pharmacies are expected to benefit from the current awareness and education campaign. These test results revealed a lack of knowledge, indicating the need for education to dispel myths and highlight the serious risks associated with type 2 diabetes mellitus. The study discovered that participants’ understanding of diabetes disease prevention through lifestyle and dietary changes was inadequate. A specialist-led educational program may increase knowledge among visitors who participate. In order to prevent the spread of diabetes, more campaigns and health-promoting prevention educational activities are required.

## 1. Introduction

Diabetes mellitus (D.M.) is a chronic hyperglycemic condition caused by a group of metabolic disorders marked by a lack of insulin production, insulin activity, or both. Diabetes-induced hyperglycemia causes various long-term damage to many organs and tissues, particularly the eyes, kidneys, blood vessels, heart, and neuronal tissue.[1] It has become one of the top ten causes of death worldwide, with a 70% increase since 2000.[2] Diabetes is classified into four general categories: type I diabetes, type II diabetes, gestational diabetes, and a particular type caused by other factors. However, type II diabetes is the most common, accounting for 90–95% of all cases [3], which, in addition to demographic determinants, is significantly influenced by modifiable risk factors such as obesity [4], physical inactivity[5], smoking[6], and dietary factors [7].

According to the International Diabetes Federation (IDF), in its tenth edition of the diabetes atlas, 2021, more than one out of every ten adults worldwide now have diabetes, and this number will continue to rise rapidly in the future [8]. Furthermore, diabetes is increasing exponentially worldwide, particularly in Africa and the Middle East, with a 156% increase in Africa and a 110% increase in the Middle East and North Africa by 2045, according to the International Diabetes Federation. The problem is that approximately half of people with diabetes are unaware that they have the disease. Because there are few signs or symptoms in the early stages of diabetes, unnoticed symptoms will persist until diabetic complications appear just before D.M. is diagnosed [9]. The reason for this rapid rise in diabetes rates is that the Middle Eastern population has one of the highest obesity rates in the world, which is known to be the primary risk factor for D.M. and its complications [10, 11]. Highlighting the importance of diabetes screening in developing countries to detect undiagnosed D.M. patients or those with high-risk factors for developing D.M. Furthermore, this will be highly beneficial in establishing a prevention program for undiagnosed D.M. individuals in order to prevent this disease [12, 13]. Jordan, a country in the Middle East, has one of the highest rates of obesity and smoking in the region. Surprisingly, significant D.M. mortality rates have not emerged in the Jordanian population [14]. Ajlouni et al. (2008) reported that the age-standardized prevalence of impaired fasting glucose (IFG) and diabetes was 7.8% and 17.1%, respectively, in Jordanians, with no statistically significant difference between men and women. [15]. The American Diabetes Association (ADA) developed the diabetes risk test, a short questionnaire, to aid in detecting both pre- and type 2 diabetes. It consists of seven questions with a score range of 0 to 11 on sex, age, hypertension, physical activity, gestational diabetes, family history of diabetes, and obesity (using a height-weight chart to reference body mass index (BMI)). This score assesses the risks of undiagnosed pre-DM patients and identifies undiagnosed D.M. patients at high risk [16].

This study aims to determine Jordanians’ awareness of type 2 diabetes among those who visit community pharmacies in Amman, Jordan, as well as clarify the role of community pharmacies in T2DM screening.

## 2. Materials and methods

### 2.1 Study Design

This study was conducted from September to December 2021 in the Amman, Jordan, community pharmacies. The study was approved by the Institutional Review Board (IBR) of the University of Petra (Decision number: 202109009), and informed consent was obtained from all the participants. All participants were reminded that their participation was entirely voluntary and that they could opt-out at any time.

A structured questionnaire was used to collect and compare demographic and knowledge data. While these sheets were distributed in community pharmacies, customers were interviewed by our trained pharmacist volunteers.

After ensuring that all participants answered all questions, all sheets were collected from customers. The ADA diabetes risk scorecard test was used to calculate the individual risk of developing type 2 diabetes mellitus. Blood sugar testing with a finger prick was performed on patients who scored more than 5 (high-risk individuals).

### 2.2 Sample size

For the Amman population of approximately four million people, the sample size was calculated using Slovin’s formula [17], a 95% confidence interval (CI), and a 0.05 margin of error. As a result, a sample size of 267 people was required to achieve the required confidence interval.

### 2.3 Subjects

The current study included 305 adult customers who visited community pharmacies in Amman, Jordan.

### 2.4 American diabetes association risk scorecard

Seven questions were developed to determine the likelihood that the general public will develop diabetes in the future [18]. The American Diabetes Association previously published the current study’s questions.

### 2.5 Instrument Development

A questionnaire was created and modified using information from the literature. The survey asked about knowledge and demographics.

The survey’s knowledge section included 14 questions that tested respondents’ general knowledge of the risks of type 2 diabetes mellitus. Each item has a checkbox to select. One item was included to ensure that participants could differentiate between inter-sex susceptibility to type 2 diabetes mellitus. Females were more likely than males to develop diabetes mellitus. The sum of correct answers was used to calculate scores, with 13 being the highest possible score value.

A cut-off value for knowledge level was used to distinguish between poor and high knowledge. If a participant scored seven or higher, he or she was considered knowledgeable, whereas if they scored less than 7, they were considered less knowledgeable. These values were statistically tested for associations among all variables. The Pearson chi-squared test is used for categorical variables (including knowledge and demographics).

The validity and reliability of the questionnaires were not tested. All statistical analyses were carried out using the Statistical Package for Social Science, with a significance set at *p* < 0.05 (version 25.0, SPSS, Chicago, IL, USA). All completed questionnaires were reviewed, and frequencies were calculated following the results. The factors influencing the knowledge of the Jordanian participants visiting community pharmacies were investigated using Multivariate logistic regression.

## 3. Results

### 3.1 Demographics of participants

Out of 315 completed questionnaires (response rate = 97%), ten forms (3.2%) were excluded from the study because respondents declined to continue the interview. Only 305 people with an ADA risk score of 5 or higher took the survey and had their blood drawn. We identified 132 individuals with risk scores of five and above (~44%). Male participants made up 54.1% (n = 165) of the total, while female participants made up 45.9% (n = 140). Participants under 40 made up (n = 115, 37.7%), while those over 40 made up (n = 190, 62.3%). The capital, Amman, received 96% of the location responses. Almost half of the individuals (n = 151, 49.5%) had diplomas or lower, and the other half (n = 154, 50.5%) had degrees higher than diplomas (i.e., bachelor’s, master’s, and PhD). When asked, we discovered that most people (n = 256, 83%) did not have a medical-related degree, and only a small percentage of pharmacy visitors (n = 49, 16.1%) did. In terms of income, nearly 40% of participants refused to reveal their earnings.

The monthly income salaries were almost evenly distributed, except for those with more than 750 JDs and less than 1000 JDs, where the percentage was 3.3% (n = 10). Only one-third of participants (n = 97, 31%) had insurance, while nearly two-thirds (n = 208, 62%) did not. Most attendees were married (n = 214, 70.2%). Smokers and non-smokers made up nearly equal percentages; smokers made up (n = 138, 45%), while non-smokers made up (n = 148, 48%). Most participants (n = 177, 58.0%) are unaware that they have chronic diseases, while the remainder (n = 128, 42.0%) are aware that they do. Table 1 summarizes the patient demographic information.

**Table 1.** The demographics of the participants (n=305).

| Questions | Sample n(%) |
| --- | --- |
| **Gender** |  |
| • Female | 140 (45.9) |
| • Male | 165 (54.1) |
| **Age (years)** |  |
| • ≤ 40 | 115 (37.7) |
| • > 40 | 190 (62.3) |
| **Education level** |  |
| • Diploma or less | 151 (49.5) |
| • Higher than diploma (bachelor, master or PhD.) | 154 (50.5) |
| **Medical major degree** |  |
| • Not having a medical-related degree | 256<br>(83.9) |
| • Having a medical-related degree | 49 (16.1) |
| **Income (monthly salary in Jordanian Dinar: J.D.)** |  |
| • Less than 250 JDs | 25 (8.2) |
| • 251-500 JDs | 81 (26.6) |
| • 501-750 | 31 (10.2) |
| • 751-1000 | 10 (3.3) |
| • More than 1000 | 39 (12.8) |
| • Not willing to disclose my salary | 119<br>(39.0) |
| **Existence of health insurance** |  |
| • Not having medical insurance | 208<br>(68.2) |
| • Having medical insurance | 97 (31.8) |
| **Social status** |  |
| • Married | 214<br>(70.2) |
| • Single | 71 (23.3) |
| • Divorced/widowed | 20 (6.6) |
| **Smoking habits** |  |
| • Smoker | 138<br>(45.2) |
| • Quit smoking | 16 (5.2) |
| • Non-smoker | 148 (48.5) |
| **Existence of chronic diseases** |  |
| • Not having chronic diseases | 177<br>(58.0) |
| • Having chronic diseases | 128 (42.0) |

**Table 2.** The overall logistic regression analysis (Multivariate) to evaluate predictors related to participants’ knowledge.

| Independent factors | Multivariate |  |
| --- | --- | --- |
|  | Beta | *p*-value |
| **Age** |  |  |
| • ≤ 40 years |  |  |
| • > 40 years | 0.062 | 0.326 |
| **Sex** |  |  |
| • Female |  |  |
| • Male | -0.038 | 0.531 |
| **Education level** |  |  |
| • Diploma or less |  |  |
| • Higher than diploma (bachelor, master or PhD.) | 0.059 | 0.335 |
| **Medical Degree** |  |  |
| • Not having a medical-related degree |  |  |
| • Having a medical-related degree | 0.198 | **0.017** |
| **Salary** |  |  |
| • ≤500 |  |  |
| • >500 | -0.037 | **0.018** |
| **Smoking habits** |  |  |
| • Smoker |  |  |
| • Non-smoking | 0.047 | 0.113 |

When all variables were tested according to knowledge level using a cut-off point of 7, the results revealed that only major medical participants had significantly more knowledge than non-medical major participants (p < 0.012).

### 3.2 Prediabetics and diabetics were identified due to the community pharmacy screening program

After testing all high-risk patients with an ADA score of 5 or higher, we discovered that 21 people out of 305, or 7.3% of all participants, had random blood sugar levels higher than 140 mg/dL but lower than 200 mg/dL. We also found five people with dangerously undiagnosed blood sugar levels above 200 mg/dL (range: 212-226 mg/dL). Such findings demonstrate how essential community pharmacies and pharmacists are in detecting type 2 diabetes mellitus.

The multivariate logistic regression results revealed a significant positive correlation (p <0.05) between participants’ knowledge and the following variables: type of college degree of participants and salary of participants, Table 4.

## 4. Discussion

Results show a significant (*p* < 0.012) correlation between the educational type of the study (medical) and the level of knowledge of risk factors for type 2 diabetes mellitus. Knowledge scores were nearly evenly distributed across subjects (52.2% of participants scored seven or more, while 47.8% scored less than seven). These findings revealed that knowledge was below average, indicating a need for educational intervention to clarify misconceptions and highlight the severe risks of type 2 diabetes mellitus. According to the participants’ knowledge scores (n = 175), 57.4% scored less than 7, and (n = 130), 42.6% scored more than 7. These test results revealed a lack of knowledge, indicating the need for educational intervention to dispel myths and highlight the severe risks associated

Even though the majority (n = 82, 68.9%) had higher degrees (master’s and PhD), those levels did not improve their knowledge. No significance was found when categorical variables (including knowledge and demographics) were tested. In line with our findings, Pan et al. (2015) reported in a Brazilian study that diabetic elderly Brazilians with low education are nearly eight times more likely to have poor knowledge of D.M. disease than those with higher education [6]. Furthermore, Hu et al.[8] previously demonstrated these relationships in a Chinese study published in 2022. A similar Swedish study discovered that having a high educational level is generally a risk factor for diabetes [9]. This study discovered that participant knowledge level is unaffected by income or smoking status. There was no correlation between these factors (income and smoking) and knowledge of the risks of developing D.M. disease after testing them. In contrast to our findings, other researchers discovered that low income caused many diabetic patients to experience diabetes distress, poor adherence to D.M. drugs, and uncontrolled blood sugar levels [10, 11]. The authors showed that low-income areas had more D.M. and obesity [12].

Almost half of our participants were smokers (n = 138, 45.2%); this reflects very high smoking rates agreeing with the previous finding, which showed a high prevalence of smoking in Jordan. [19–22]. In a recent report, the kingdom of Jordan was classified as the sixth internationally; smoking prevalence was almost 70% between males and approximately 10% among females [23]. This risk factor was recently added to the list of risk factors for diabetes mellitus. [24] another report confirmed this causal relationship between smoking and especially type 2 diabetes mellitus from Japan.[25] Another significant report highlights that quitting smoking is crucial not just for preventing macrovascular problems in diabetes but also for minimizing microvascular damage and may help with glycemic control in this condition. [26] Confirming relations between smoking and D.M. complications[27].

Nearly 62 per cent of our cohort participants were over 40 years old (n = 190, 62.3%); this is considered the starting risk for developing type 2 diabetes mellitus. [28] We identified 132 individuals with risk scores of five and above (~44%); This showed high risks among random community pharmacy visitors and agreed with national reports that confirmed a high rate of prediabetics among Jordanians.[29]

## 5. Limitations of the study

The short duration of this study was a significant flaw. There was no follow-up monitoring to see if improved knowledge and daily activities affected long-term outcomes. Furthermore, because we only screened individuals in the Amman governorate, we cannot generalize our findings to the rest of Jordan’s cities.

## 6. Conclusions

This study discovered that participants’ understanding of D.M. disease prevention through lifestyle and dietary changes was inadequate. Additional educational initiatives by a specialist may raise participants’ knowledge levels. Additional campaigns and health-promoting prevention education programs are required to continue preventing D.M. disease.

## Acknowledgements

We want to offer our heartfelt gratitude to the Faculty of Pharmacy and Medical Sciences at the University of Petra for allowing us to complete this research successfully. The authors would also like to thank the Deanship of Scientific Research and Graduate Studies for their invaluable assistance. Furthermore, we would like to acknowledge the role of Professor Hisham Al Matubsi’s outstanding contributions in editing and revising the final draft of the manuscript.

## Contribution of Authors

We declare that this work was done by the authors named in this article, and the authors will bear all liabilities about claims relating to the content of this article.

